# Divergent LD-transpeptidase-independent effects of peptidoglycan carboxypeptidases on intrinsic β-lactam and vancomycin resistance

**DOI:** 10.1101/2022.04.04.487088

**Authors:** Si Hyoung Park, Umji Choi, Su-Hyun Ryu, Han Byeol Lee, Jin-Won Lee, Chang-Ro Lee

**Author notes:** These authors have contributed equally to this work (Si Hyoung Park, Umji Choi). **For correspondence:** Chang-Ro Lee.

## Abstract

Vancomycin and β-lactams are clinically important antibiotics that inhibit the formation of peptidoglycan cross-links, but their binding targets are different. The binding target of vancomycin is D-alanine-D-alanine (D-Ala-D-Ala), whereas that of β-lactam is penicillin-binding proteins (PBPs). In this study, we revealed the divergent effects of peptidoglycan (PG) carboxypeptidases on vancomycin and β-lactam resistance in *Escherichia coli* and *Bacillus subtilis*. The deletion of PG carboxypeptidases induced sensitivity to most β-lactams, whereas it induced strong resistance toward vancomycin. Notably, both of two phenotypes did not have strong association with LD-transpeptidases, which are necessary for the formation of PG 3-3 cross-links and covalent bonds between PG and an Lpp outer membrane (OM) lipoprotein. Vancomycin resistance was induced by increased amount of decoy D-Ala-D-Ala residues within PG, whereas β-lactam sensitivity was associated with physical interactions between PG carboxypeptidase and PBPs. The presence of OM permeability barrier strongly strengthened vancomycin resistance, but it significantly weakened β-lactam sensitivity. Collectively, our results revealed two distinct LD-transpeptidase-independent functions of PG carboxypeptidases, which involved inverse modulation of bacterial resistance to clinically important antibiotics, β-lactams and vancomycin, and presented evidence for a link between PG carboxypeptidase and PBPs.

**IMPORTANCE:** Bacterial peptidoglycan (PG) hydrolases play important roles in various aspects of bacterial physiology, including cytokinesis, PG synthesis, quality control of PG, PG recycling, and stress adaptation. Of all the PG hydrolases, the role of PG carboxypeptidases is poorly understood, especially regarding their impacts on antibiotic resistance. To date, most studies on PG carboxypeptidases are focused on LD-transpeptidase-related roles. We have revealed two distinct LD-transpeptidase-independent functions of PG carboxypeptidases with respect to antibiotic resistance. The deletion of PG carboxypeptidases led to sensitivity to most β-lactams, while it caused strong resistance to vancomycin. The underlying molecular mechanisms of two phenotypes were not associated with LD-transpeptidases. Therefore, our study provides novel insights into the roles of PG carboxypeptidases in the regulation of antibiotic resistance and a potential clue for the development of a drug to improve the clinical efficacy of β-lactam antibiotics.

**One sentence summary:** Effect of peptidoglycan carboxypeptidase on antibiotic

## INTRODUCTION

Peptidoglycan (PG) is a pivotal mesh-like structure that is necessary for shape maintenance and protection from turgor pressure-mediated lysis (1). PG is composed of linear amino sugar polymers and short peptide chains. The linear glycan strand consists of alternating β-1,4-linked two amino sugars, N-acetylglucosamine (GlcNAc) and N-acetylmuramic acid (MurNAc). The short peptide chain is attached to the D-lactoyl moiety of each MurNAc. In many bacteria, including *Escherichia coli*, the short peptide chain is composed of three D-amino acids, one L-amino acid, and one biosynthetic precursor of L-lysine (L-alanine, D-glutamate, *meso*-diaminopimelate, D-alanine, and D-alanine). The integrity of PG is strengthened by covalent cross-links between peptide chains. In *E. coli*, the cross-link is predominantly (more than 90%) formed between the fourth D-alanine of the donor peptide and the third *meso*-diaminopimelate (*meso*-DAP) of the acceptor peptide (4-3 cross-links) and rarely (approximately 3~10%) between the third *meso*-DAP and the third *meso*-DAP (3-3 cross-links) (2).

Bacteria have many PG hydrolases that play a role in PG synthesis and degradation (3). Lytic transglycosylases hydrolyze the β-1,4-glycosidic bond between GlcNAc and MurNAc, whereas PG amidases and peptidases cleave the cross-linked peptides. PG amidases cleave the lactylamide bond between MurNAc and the peptide chain, and promote cell separation during cytokinesis (4, 5). PG peptidases are classified into two subgroups, endopeptidases that hydrolyze the cross-linked bridges and carboxypeptidases that cleave the fifth D-alanine of the peptide chain (3). *E. coli* has seven PG endopeptidases, which seem to play distinct roles in cross-link cleavage for PG synthesis (6), and seven PG carboxypeptidases (3). After the removal of the fifth D-alanine by PG carboxypeptidases, the third *meso*-DAP of the PG tetrapeptide can be covalently cross-linked to the third *meso*-DAP of adjacent peptide chains by some LD-transpeptidases (LdtD and LdtE) or to the lysine residue of an abundant outer membrane (OM) lipoprotein Lpp (Braun’s lipoprotein) by other LD-transpeptidases (LdtA, LdtB, and LdtC) (7–9). The covalent cross-links between the peptide chain of PG and Lpp provide a tight connection between PG and OM, which increases the envelope integrity (10). However, the physiological significance of 3-3 cross-links remains unclear. A recent study showed that the 3-3 cross-link might strengthen the PG under the defect of OM synthesis (7).

One of the evident physiological roles of 3-3 cross-links is β-lactam resistance (2). All β-lactams inhibit DD-transpeptidases (also known as penicillin-binding protein, PBP) that catalyze the formation of 4-3 cross-links, but most β-lactams, except carbapenems, do not inhibit LD-transpeptidases (11). Therefore, the *E. coli* mutant with increased 3-3 cross-links demonstrated elevated resistance to β-lactams (2). In addition, several pathogenic bacteria, including *Mycobacterium tuberculosis* and *Clostridium difficile*, in which 3-3 cross-links are highly abundant (more than 60%), were highly resistant to β-lactams (11, 12). Furthermore, the *E. coli* mutant defective for DacA (also known as PBP5) exhibited increased sensitivity to β-lactams (13–15). Given that the activity of PG carboxypeptidases is necessary for the formation of 3-3 cross-links, the decrease in the proportion of 3-3 cross-links should ideally result in increased sensitivity of the *dacA* mutant to β-lactams. However, this may not occur due to low intracellular proportion (approximately 3–10%) of the 3-3 cross-link.

In this study, we presented two divergent phenotypes of the *dacA* mutant, β-lactam sensitivity and vancomycin resistance. Unexpectedly, the two phenotypes were not mainly associated with both of 3-3 cross-link formation and the Lpp-PG attachment catalyzed by LD-transpeptidases. Vancomycin resistance was induced by increase in decoy D-Ala-D-Ala residues, while β-lactam sensitivity was associated with physical interactions between DacA and PBPs. The DacA(ΔC) without its C-terminal membrane-anchoring domain showed completely distinct effects on the two phenotypes of the *dacA* mutant. Likewise, the presence of OM permeability barrier also differentially affected the two phenotypes. Our study revealed two distinct LD-transpeptidase-independent molecular mechanisms of PG carboxypeptidase-mediated antibiotic resistance and showed a novel functional relationship between PG carboxypeptidase and PBPs.

## RESULTS

### Deletion of PG carboxypeptidases causes vancomycin resistance in *E. coli*

To analyze the physiological functions of PG carboxypeptidases, we examined several envelope stress-related phenotypes of PG carboxypeptidase-deleted strains. Notably, phenotypes were detected only in the *dacA* mutant (Fig. S1). Among the identified phenotypes, we focused on vancomycin resistance. The *dacA* mutant exhibited strong vancomycin resistance, whereas the growth of other PG carboxypeptidase-deleted mutants was almost completely comparable to that of the wild-type strain in the presence of vancomycin (Fig. 1A). To assess whether the PG carboxypeptidase activity of DacA is involved in vancomycin resistance, we constructed DacA(S73G) mutant in which the serine residue of the active site was substituted with glycine, which resulted in the complete loss of PG carboxypeptidase activity (16). Vancomycin resistance of the *dacA* mutant was complemented by the ectopic expression of the wild-type DacA protein, but not DacA(S73G) mutant protein (Fig. 1B), indicating that vancomycin resistance is associated with the PG carboxypeptidase activity of DacA.

**FIG 1.**
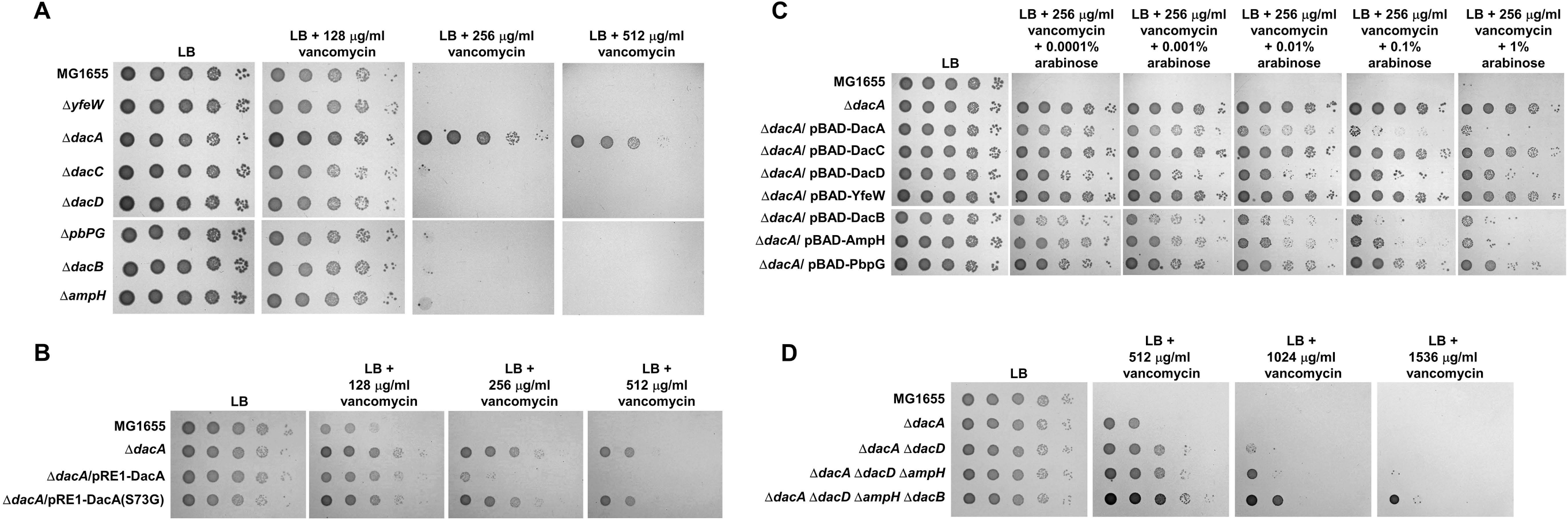
The loss of PG carboxypeptidase activities induces vancomycin resistance. (A) Vancomycin resistance of the *dacA* mutant. The wild-type and indicated mutant cells were serially diluted from 10^8^ to 10^4^ cells/ml in 10-fold steps and spotted onto an LB plate and LB plates containing indicated vancomycin concentrations. (B) Complementation of vancomycin resistance of the *dacA* mutant. The cells of the indicated strains were serially diluted from 10^8^ to 10^4^ cells/ml in 10-fold steps and spotted onto an LB plate and LB plates containing the indicated vancomycin concentrations. (C) Complementation of vancomycin resistance of the *dacA* mutant by other PG carboxypeptidases. The cells of the indicated strains were serially diluted from 10^8^ to 10^4^ cells/ml in 10-fold steps and spotted onto an LB plate and LB plates containing 256 μg/ml vancomycin and the indicated concentrations of arabinose. (D) Enhancement of vancomycin resistance by additional deletions of other PG carboxypeptidases. The wild-type and indicated mutant cells were serially diluted from 10^8^ to 10^4^ cells/ml in 10-fold steps and spotted onto an LB plate and LB plates containing indicated vancomycin concentrations.

Because the PG carboxypeptidase activity of DacA is important in vancomycin resistance, we determined whether the expression of other proteins with PG carboxypeptidase activity can influence vancomycin resistance of the *dacA* mutant. When PG carboxypeptidases were expressed using the arabinose-inducible promoter, vancomycin resistance of the *dacA* mutant was reduced by the expression of DacB, AmpH, DacD, and DacA (Fig. 1C). These results imply that additional deletion of other PG carboxypeptidases in the *dacA* mutant might increase vancomycin resistance. To test this assumption, we constructed the *dacA dacD* double, *dacA dacD ampH* triple, and *dacA dacD ampH dacB* quaternary mutants. Expectedly, vancomycin resistance of the *dacA* mutant gradually increased along with increase in the number of deleted genes (Fig. 1D). These results demonstrate that the loss of PG carboxypeptidases is associated with vancomycin resistance and DacA plays a major role in this phenotype.

### Deletion of DacA results in β-lactam sensitivity

Some previous studies have demonstrated that the *dacA* mutant is sensitive to β-lactams (13–15). We also observed ampicillin sensitivity of the *dacA* mutant, and as observed for vancomycin resistance, this phenotype was detected only in the *dacA* mutant (Fig. 2A). The ampicillin sensitivity was also associated with the PG carboxypeptidase activity of DacA (Fig. 2B). Because vancomycin resistance was complemented by the expression of other PG carboxypeptidases, we examined whether the ampicillin sensitivity is also restored by the expression of other PG carboxypeptidases. Besides DacA, several other PG carboxypeptidases could partially complement the ampicillin sensitivity of the *dacA* mutant (Fig. 2C). The additional deletions of PG carboxypeptidases in the *dacA* mutant strengthened ampicillin sensitivity (Fig. 2D), although the effect was not as much as that for vancomycin resistance. In summary, these results indicate that PG carboxypeptidases are involved in ampicillin sensitivity and DacA plays a pivotal role in this phenotype.

**FIG 2.**
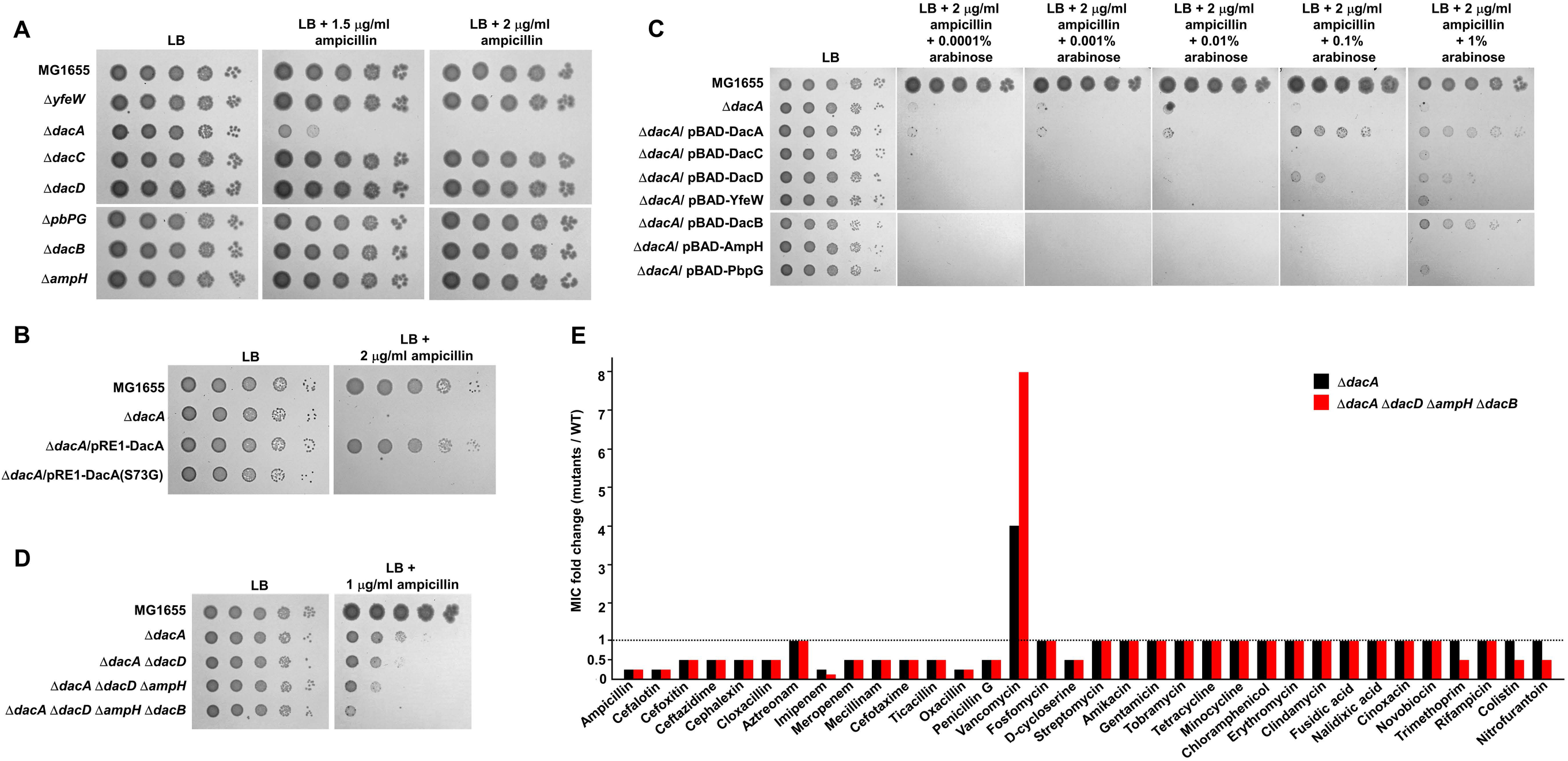
The deletion of the *dacA* gene induces β-lactam sensitivity. (A) Ampicillin sensitivity of the *dacA* mutant. The wild-type and indicated mutant cells were serially diluted from 10^8^ to 10^4^ cells/ml in 10-fold steps and spotted onto an LB plate and LB plates containing the indicated ampicillin concentrations. (B) Complementation of ampicillin sensitivity of the *dacA* mutant. The cells of the indicated strains were serially diluted from 10^8^ to 10^4^ cells/ml in 10-fold steps and spotted onto an LB plate and an LB plate containing 2 μg/ml ampicillin. (C) Complementation of ampicillin sensitivity of the *dacA* mutant by other PG carboxypeptidases. The cells of the indicated strains were serially diluted from 10^8^ to 10^4^ cells/ml in 10-fold steps and spotted onto an LB plate and LB plates containing 2 μg/ml ampicillin and the indicated concentrations of arabinose. (D) The effect of additional deletions of other PG carboxypeptidases on ampicillin sensitivity of the *dacA* mutant. The wild-type and indicated mutant cells were serially diluted from 10^8^ to 10^4^ cells/ml in 10-fold steps and spotted onto an LB plate and an LB plate containing 2 μg/ml ampicillin. (E) The effect of PG carboxypeptidases on the MICs of antibiotics. The MICs of various antibiotics were measured against the wild-type or indicated mutant strains. The relative MIC values for the mutant cells compared to those for the wild-type cells are shown.

The inverse phenotypes of the *dacA* mutant for vancomycin and ampicillin susceptibility prompted us to investigate how susceptibility to other antibiotics is also affected by the *dacA* deletion. We examined the minimal inhibitory concentrations (MICs) of diverse antibiotics against the wild-type and *dacA* mutant strains. The MIC of ampicillin was 4-fold lower for the *dacA* mutant than for the wild-type strain; moreover, the MICs of most β-lactams were also 2-fold or 4-fold lower for the *dacA* mutant (Fig. 2E). Besides β-lactams, the MIC of D-cycloserine was also 2-fold lower for the *dacA* mutant than for the wild-type strain. The only antibiotic whose MIC is higher for the *dacA* mutant than for the wild-type strain was vancomycin (4-fold higher) (Fig. 2E). It is interesting that the *dacA* mutant exhibited divergent phenotypes to two antibiotics, β-lactam and vancomycin, inhibiting PG cross-linking. Furthermore, additional deletions of PG carboxypeptidases in the *dacA* mutant differentially affected the MICs of two antibiotics. The MIC of vancomycin was 2-fold higher in the *dacA dacD ampH dacB* quaternary mutant than in the *dacA* mutant, while the MICs of most β-lactams were the same in the two strains (Fig. 2E). In summary, these results indicate that DacA divergently modulates the MICs of vancomycin and β-lactams. Other PG carboxypeptidases also mildly influence the MIC of vancomycin, but their effects on β-lactams are very weak, implying that the underlying mechanisms of divergent phenotypes of the *dacA* mutant against two antibiotics are distinct.

### Neither vancomycin resistance nor β-lactam sensitivity of the *dacA* mutant has a strong association with LD-transpeptidases

After the removal of the fifth D-alanine by PG carboxypeptidases, the PG tetrapeptide containing four amino acids can form the 3-3 cross-links by LD-transpeptidases (LdtD and LdtE) (2, 7, 9). LD-transpeptidases are structurally unrelated to PBPs (17); thus, they are not inhibited by most β-lactam antibiotics (11). Although these 3-3 cross-links account for only 3–10% of the entire cross-links in *E. coli* (18), a mutant strain having increased proportion of 3-3 cross-links showed β-lactam resistance, owing to the reduction in PBP dependency (2). To assess whether the 3-3 cross-links are involved in β-lactam sensitivity of the *dacA* mutant, we examined the ampicillin susceptibility of the *ldtD ldtE* double mutant. Notably, the ampicillin sensitivity of the *ldtD ldtE* double mutant was comparable to that of the wild-type strain (Fig. 3A). After the removal of the fifth D-alanine by PG carboxypeptidases, the PG tetrapeptide can also be covalently cross-linked to Lpp by other LD-transpeptidases (LdtA, LdtB, and LdtC) (7). We checked the effect of LdtABC or Lpp on ampicillin sensitivity. The susceptibility of the *ldtABC* or *lpp* mutant to ampicillin was also the same as that of the wild-type stain (Fig. 3A). Additionally, the *ldtDE lpp* mutant defective for both the 3-3 cross-link and the attachment of PG to Lpp also exhibited the susceptibility comparable to the wild-type strain (Fig. 3A). To assess whether the same result can be obtained for other β-lactam antibiotics, we determined the MICs of various β-lactams against the *ldtDE lpp* mutant. Expectedly, for the *ldtDE lpp* mutant, there were no changes in the MICs of all β-lactams tested, compared to those observed for the wild-type (Fig. 3B). These results strongly suggest that the β-lactam sensitivity of the *dacA* mutant is not related to LD-transpeptidases.

**FIG 3.**
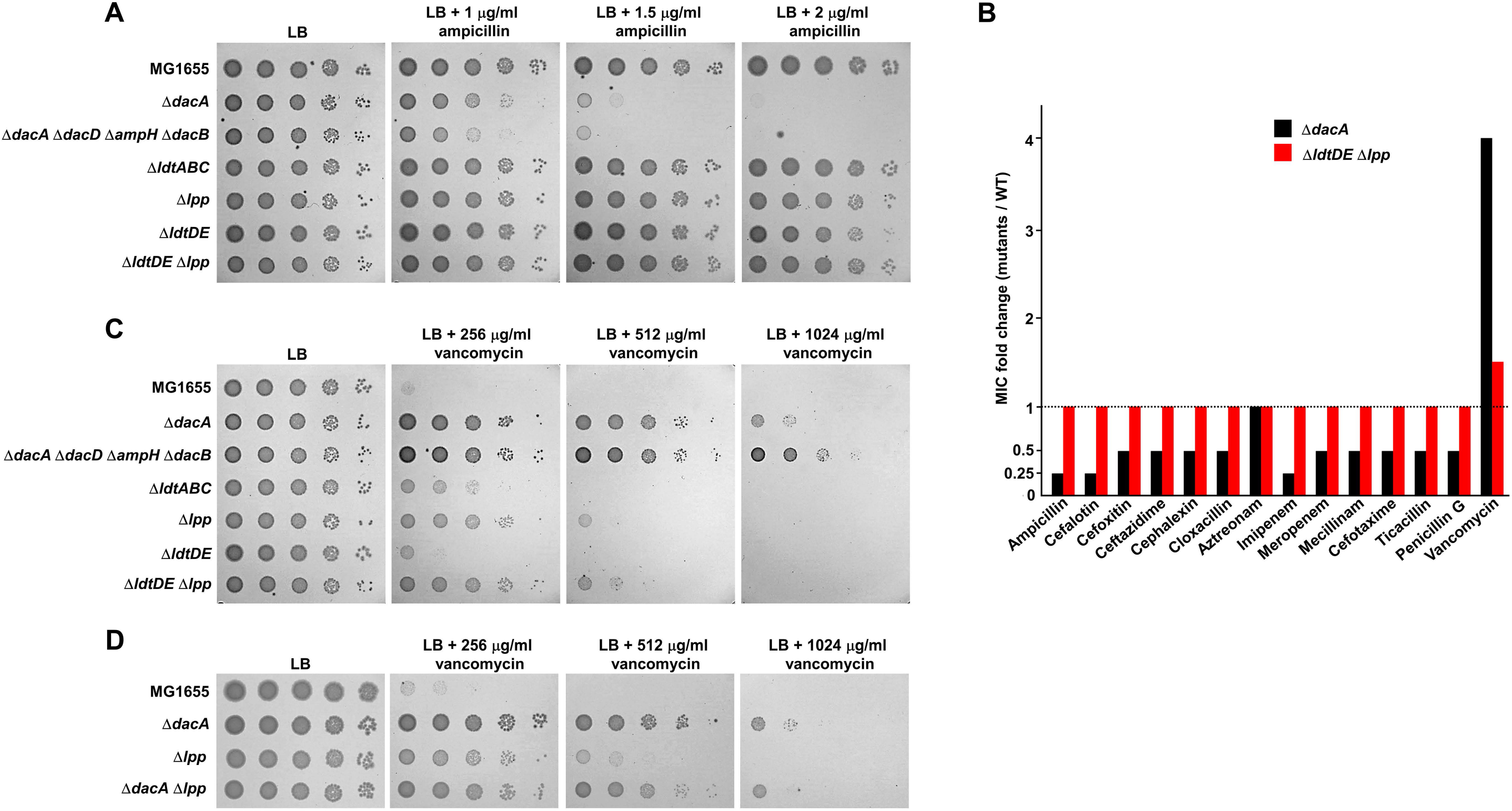
Phenotypes of the *dacA* mutant are not mainly associated with LD-transpeptidases. (A) The effect of LD-transpeptidases on ampicillin susceptibility. The wild-type and indicated mutant cells were serially diluted from 10^8^ to 10^4^ cells/ml in 10-fold steps and spotted onto an LB plate and LB plates containing the indicated ampicillin concentrations. (B) The effect of LD-transpeptidases on the MICs of β-lactams. The MICs of various antibiotics were measured against the wild-type or indicated mutant strains. The relative MIC values for the mutant cells compared to those for the wild-type cells are shown. (C) The effect of LD-transpeptidases on vancomycin resistance. The wild-type and indicated mutant cells were serially diluted from 10^8^ to 10^4^ cells/ml in 10-fold steps and spotted onto an LB plate and LB plates containing indicated vancomycin concentrations. (D) The effect of additional deletion of Lpp on vancomycin resistance of the *dacA* mutant. The wild-type and indicated mutant cells were serially diluted from 10^8^ to 10^4^ cells/ml in 10-fold steps and spotted onto an LB plate and LB plates containing the indicated vancomycin concentrations.

Next, we analyzed the relationship between vancomycin resistance and LD-transpeptidases. The vancomycin susceptibility of the *ldtDE* mutant was comparable to that of the wild-type cells, but both the *ldtABC* and *lpp* mutants were slightly more resistant to vancomycin than the wild-type cells, and the vancomycin resistance of the *ldtDE lpp* mutant was comparable to that of the *lpp* mutant (Fig. 3C). The MIC of vancomycin was 1.5-fold higher for the *ldtDE lpp* mutant than for the wild-type strain (Fig. 3B). The vancomycin resistance of the *dacA* mutant was not strengthened by the additional deletion of Lpp (Fig. 3D). Collectively, these results imply that the loss of the cross-links between PG and Lpp partially contributes to vancomycin resistance of the *dacA* mutant. Because vancomycin resistance of the *ldtDE lpp* mutant was significantly weaker than that of the *dacA* or *dacA dacD ampH dacB* mutant, the loss of the Lpp-PG attachment may not be the main molecular basis of vancomycin resistance of the PG carboxypeptidase-defective strains.

### Increase in decoy D-Ala-D-Ala residues within PG in the *dacA* mutant results in vancomycin resistance

The fifth D-Ala of the donor pentapeptides is removed during the formation of the 4-3 cross-links by PBPs, whereas the fifth D-Ala of the acceptor pentapeptides is not removed during these reactions by PBPs (19). Because the fifth D-Ala of the acceptor pentapeptides can be removed by PG carboxypeptidases (19), the deletion of PG carboxypeptidases increases the amount of the 4-3 cross-links with D-Ala-D-Ala at the acceptor peptides (20). These D-Ala-D-Ala residues of cross-linked stem peptides can act as a decoy for the interaction with vancomycin, thus decreasing the amount of free vancomycin that inhibits the function of PBPs and consequently, increasing vancomycin resistance. Because in *E. coli*, the amount of vancomycin passing through the periplasm is very low due to the presence of OM (21), the increase in decoy D-Ala-D-Ala residues can strongly affect the MIC of vancomycin. To assess this assumption, we examined the amount of vancomycin bound to purified PG using fluorescent vancomycin derivative. The amount of bound fluorescent vancomycin was approximately 3.5-fold higher in PG isolated from the *dacA* mutant than in that from the wild-type strain (Fig. 4A). The difference highly correlated quantitatively with that in MIC (Fig. 3E) when the weak vancomycin resistance induced by the loss of the Lpp-PG attachment was considered. Therefore, these results strongly suggest that the increase in decoy D-Ala-D-Ala residues at the acceptor peptides induced by the deletion of PG carboxypeptidases causes vancomycin resistance.

**FIG 4.**
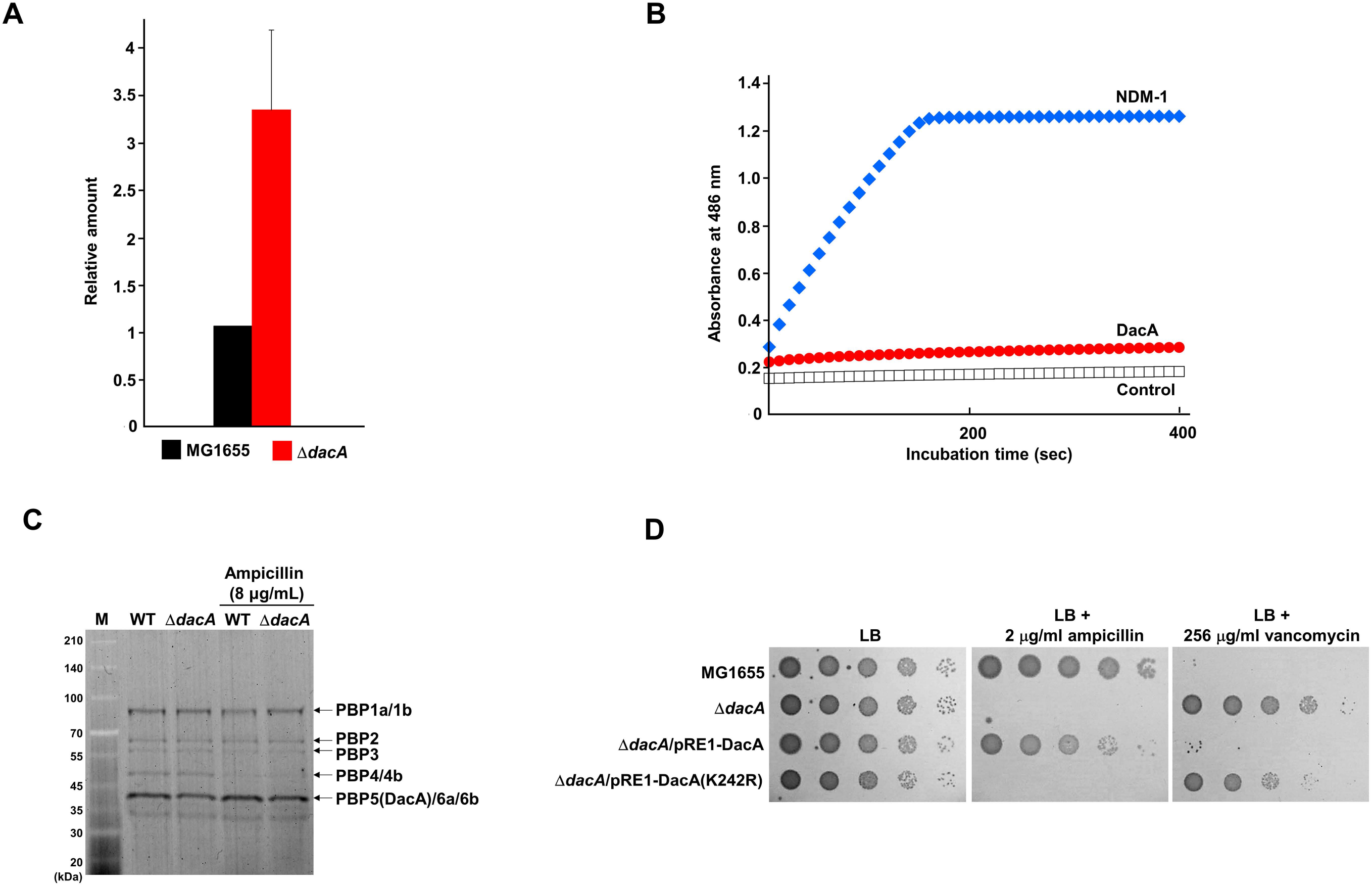
The analysis of diverse DacA-related physiological properties. (A) *In vitro* interaction between fluorescent vancomycin and purified PG. Purified PGs from the wild-type and *dacA* mutant cells were mixed with fluorescent vancomycin. The amount of vancomycin bound to PG was determined by measuring the fluorescence emission of fluorescent vancomycin at 513 nm following excitation at 490 nm. (B) Estimation of the β-lactamase activity of DacA. The β-lactamase activities of purified NDM-1 (11 μg) and DacA (160 μg) were measured using nitrocefin. (C) Representative SDS-PAGE gel image for ampicillin titration of PBPs from the wild-type and *dacA* mutant cells. The binding assay of Bocillin FL to PBPs of the wild-type and *dacA* mutant cells was performed in the absence or presence of 8 μg/ml ampicillin. (D) Complementation of the phenotypes of the *dacA* mutant by DacA(K242R). The cells of the indicated strains were serially diluted from 10^8^ to 10^4^ cells/ml in 10-fold steps and spotted onto an LB plate and an LB plate containing 2 μg/ml ampicillin or 256 μg/ml vancomycin.

### The C-terminal domain of DacA is dispensable for its PG carboxypeptidase activity, but is obligately required for β-lactam resistance

Previous reports suggested that DacA may have weak β-lactamase activity as it shares a common ancestor with β-lactamases and contains an Ω-loop-like domain similar to TEM-1 β-lactamase (19, 22). However, till date, the β-lactamase activity of purified DacA has not been reported (23). In this study, we determined whether purified DacA has the β-lactamase activity. Purified NDM-1 showed a strong nitrocefin-degrading activity, but DacA did not degrade nitrocefin, even at a condition of approximately 14-fold higher protein level (Fig. 4B), indicating that DacA has no β-lactamase activity.

Given that DacA is one of the seven class C PBP proteins with low molecular weights (24), it can act as a β-lactam decoy protein that diminishes the concentration of free β-lactams. However, a previous report showed that the binding affinities of DacA with most β-lactams were significantly lower than those of other class C PBPs (25). Nevertheless, as DacA is the most abundant PG carboxypeptidase under normal conditions (26, 27), we assessed the role of DacA as a β-lactam decoy protein. In the wild-type and *dacA* mutant cells, the level of inhibition of Bocillin FL binding to PBPs by ampicillin was examined. In this assay, the band positions of DacA, DacC (also known as PBP6a), and DacD (also known as PBP6b) proteins were indistinguishable due to their similar molecular weights. Notably, the intensity of the band corresponding to DacA, DacC, and DacD was slightly reduced by the deletion of DacA, which was consistent with a previous report (28), indicating that DacA does not exist at an excessive level. The inhibition levels of Bocillin FL binding to PBPs by ampicillin were similar between the wild-type and *dacA* mutant strains (Fig. 4C), demonstrating that the role of DacA as the β-lactam decoy protein is negligible. To confirm this finding, we constructed a DacA(K242R) mutant whose PG carboxypeptidase activity is completely lost, but binding affinity for penicillin G is increased (29). Although this mutant protein can act as the β-lactam decoy protein, it could not complement the ampicillin sensitivity of the *dacA* mutant (Fig. 4D). Collectively, these results suggest that β-lactam sensitivity of the *dacA* mutant is not associated with the role of DacA as the β-lactam decoy protein.

DacA has an inner membrane-anchoring domain at its C-terminus (from 386^th^ to 403^th^ amino acid residues), which is important for the function of DacA in morphology and its localization at the cell division site (16, 30). To analyze the effect of this domain on vancomycin resistance and β-lactam sensitivity, we constructed DacA(ΔC) mutant without the C-terminal membrane-anchoring domain. Unexpectedly, DacA(ΔC) showed significantly increased PG carboxypeptidase activity than the wild-type protein (Fig. 5A). DacA(ΔC) could completely complement vancomycin resistance of the *dacA* mutant, confirming the full enzymatic activity of DacA(ΔC) (Fig. 5B). Notably, DacA(ΔC) could not restore ampicillin sensitivity of the *dacA* mutant (Fig. 5B), despite its full enzymatic activity. These results were also confirmed by the strain defective for a chromosomal C-terminal domain of DacA, MG1655 DacA(ΔC)-Flag. DacA(ΔC) was adequately expressed in this strain (Fig. S2) and thus, vancomycin resistance was not observed (Fig. 5B), but ampicillin sensitivity was detected up to the level comparable to that in the *dacA* mutant (Fig. 5B). The MG1655 DacA(ΔC)-Flag strain completely phenocopied the *dacA* mutant in β-lactam sensitivity (Fig. 5D), strongly indicating the obligate requirement of the C-terminal domain of DacA in β-lactam resistance. Collectively, these results again confirm the distinct molecular mechanisms of the two phenotypes of the *dacA* mutant, and strongly suggest that the C-terminal domain of DacA and its enzymatic activity are strictly required for β-lactam resistance.

**FIG 5.**
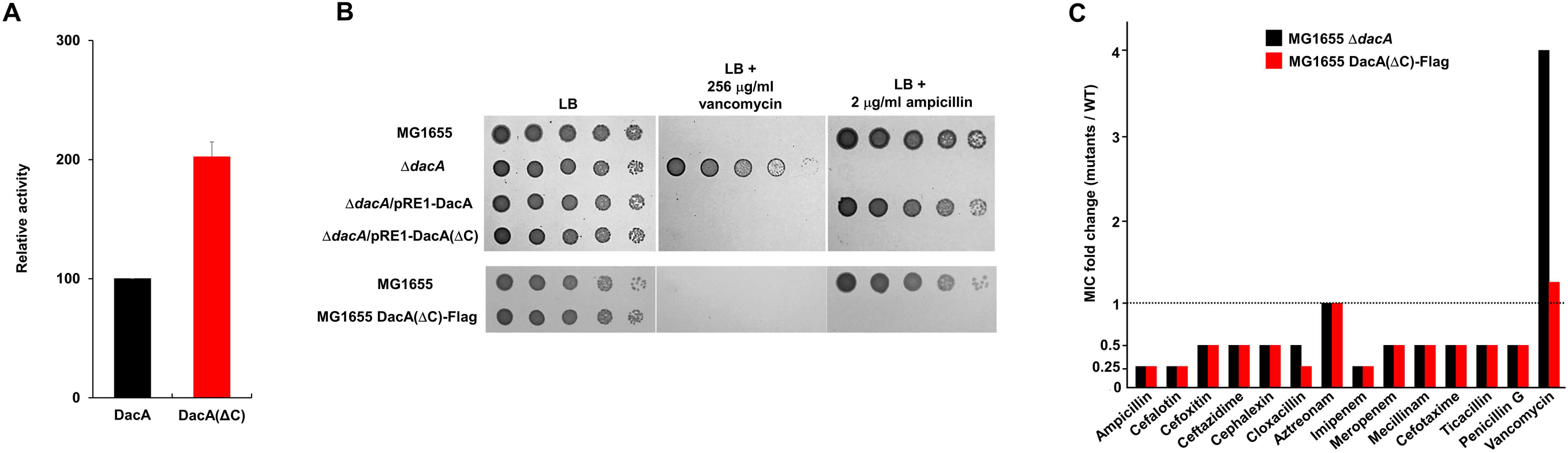
Effects of the C-terminal domain of DacA on β-lactam and vancomycin resistance. (A) Effect of the C-terminal domain of DacA on its enzymatic activity. The enzymatic activity of purified PG carboxypeptidases was measured using the substrate diacetyl-L-Lys-D-Ala-D-Ala (AcLAA). Purified proteins (1 μM) were incubated at 37°C with 50 mM Tris-HCl (pH 8.0) containing 1 mM AcLAA. Released D-Ala was estimated using Horseradish peroxidase and Amplex Red at 563 nm. (B) Effect of the C-terminal domain of DacA on β-lactam and vancomycin resistance. The cells of the indicated strains were serially diluted from 10^8^ to 10^4^ cells/ml in 10-fold steps and spotted onto an LB plate and an LB plate containing 256 μg/ml vancomycin or 2 μg/ml ampicillin. (C) The MICs of β-lactams against the MG1655 DacA(ΔC)-Flag strain. The MICs of various antibiotics were measured against the wild-type or indicated mutant strains. The relative MIC values for the mutant cells compared to those for the wild-type cells are shown.

### DacA interacts with PBPs in a C-terminal domain-dependent manner

Because the *dacA* mutant was sensitive to almost all the β-lactam antibiotics, but not to the other antibiotics with different modes of action (Fig. 2E), we analyzed the physical interaction between DacA and PBP, a binding target of β-lactam. To perform pull-down experiments, we constructed PBP-Flag strains having three Flag-tags at the C-terminus of chromosomal PBPs. All the PBP proteins, including PBP1a, PBP1b, PBP2, and PBP3, were pulled down by His-tagged DacA (Fig. 6A and Fig. S3), suggesting a physical interaction between DacA and PBPs. Notably, no PBP protein was pulled down by His-tagged DacA(ΔC) (Fig. 6B and Fig. S4), indicating that the C-terminal domain of DacA is required for these interactions. The C-terminal domain-dependent interaction of DacA with PBP was also confirmed by pull-down experiments using overexpressed PBP1a and PBP1b. Although the same overexpressed PBP1a or PBP1b supernatant was used, a significant amount of PBP1a or PBP1b was pulled down by His-tagged DacA, but not by His-tagged DacA(ΔC), even though the level of purified His-tagged DacA(ΔC) was higher than that of purified His-tagged DacA (Fig. 6C). These results suggest that DacA interacts with PBPs in a C-terminal domain-dependent manner.

**FIG 6.**
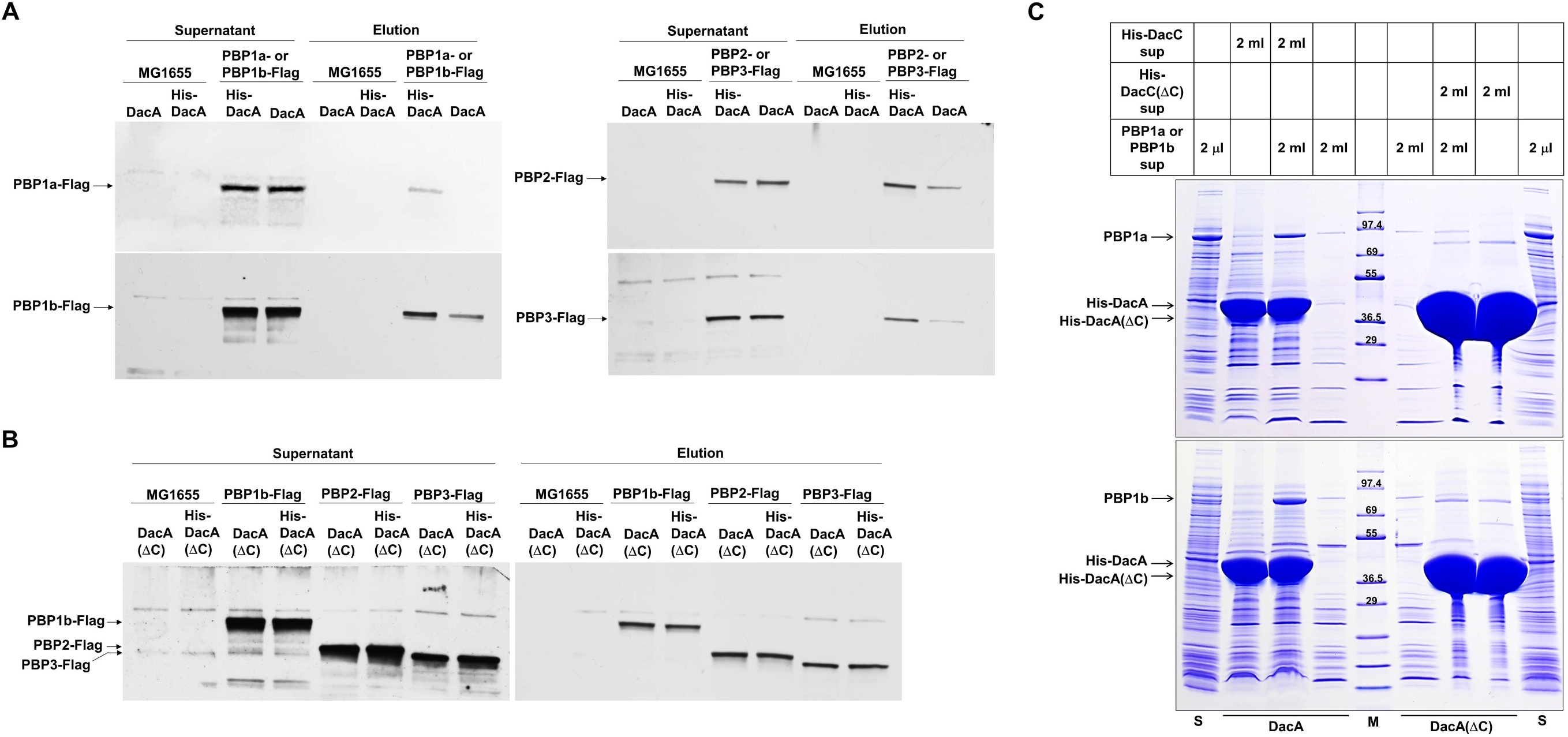
The C-terminal domain-dependent interactions of DacA with PBPs. (A) The physical interactions between DacA and PBPs. The supernatant of MG1655 cells or MG1655 cells with three Flag-tags fused to the C-terminus of the indicated chromosomal PBP was mixed with the supernatant of ER2566 cells harboring pET-based plasmid expressing His-tagged DacA or non-tagged DacA. After pull-down experiments, the amount of input (Supernatant) and output (Elution) PBPs was measured by western blot using monoclonal antibody against Flag-tag. (B) The importance of the C-terminal domain of DacA for its interaction with PBPs. The supernatant of MG1655 cells or MG1655 cells with three Flag-tags fused to the C-terminus of the indicated chromosomal PBP was mixed with the supernatant of ER2566 cells harboring pET-based plasmid expressing His-tagged DacA(ΔC) or non-tagged DacA(ΔC). After pull-down experiments, the amount of input (Supernatant) and output (Elution) PBPs was measured by western blot using monoclonal antibody against Flag-tag. (C) The C-terminal domain-dependent interaction of DacA with PBP1a or PBP1b. The supernatant of ER2566 cells harboring the pET28a plasmid expressing PBP1b was mixed with the supernatant of ER2566 cells harboring pET-based plasmid expressing His-tagged DacA or His-tagged DacA(ΔC). After pull-down experiments, eluted proteins were separated on 4–20% gradient Tris-glycine polyacrylamide gels. The supernatant (lane S) of ER2566 cells harboring the pET28a plasmid expressing PBP1a or PBP1b was also loaded onto the gel. Lane M indicates EzWayTM Protein Blue MW Marker (KOMA Biotech., Korea).

### The presence of OM permeability barrier is necessary for vancomycin resistance of the *dacA* mutant, but not for β-lactam susceptibility

The OM plays a role as a strong barrier that prevents the transport of toxic molecules, such as antibiotics (21). To determine whether the OM permeability barrier affects vancomycin resistance and β-lactam sensitivity of the *dacA* mutant, we examined the effect of *dacA* depletion in the context of the *ompA ompC ompF* triple mutant, a strain with increased OM permeability (31). The *ompA ompC ompF* triple mutant was more sensitive to vancomycin than the wild-type strain (Fig. 7A), indicating the increased passage of vancomycin in this strain. Notably, in the background of the *ompA ompC ompF* triple mutant, vancomycin resistance of the *dacA* mutant was completely absent (Fig. 7A). These results imply that the effect of the decoy D-Ala-D-Ala within PG on vancomycin resistance weakens as the periplasmic level of vancomycin increases. Interestingly, unlike vancomycin, ampicillin sensitivity of the *dacA* mutant was significantly strengthened in the background of the *ompA ompC ompF* triple mutant (Fig. 7B). This pattern was also observed for most other β-lactams apart from ampicillin (Fig. 7C), indicating that β-lactam sensitivity of the *dacA* mutant is independent of OM permeability. These data confirm that the molecular mechanisms for the two inverse phenotypes of the *dacA* mutant are distinct.

**FIG 7.**
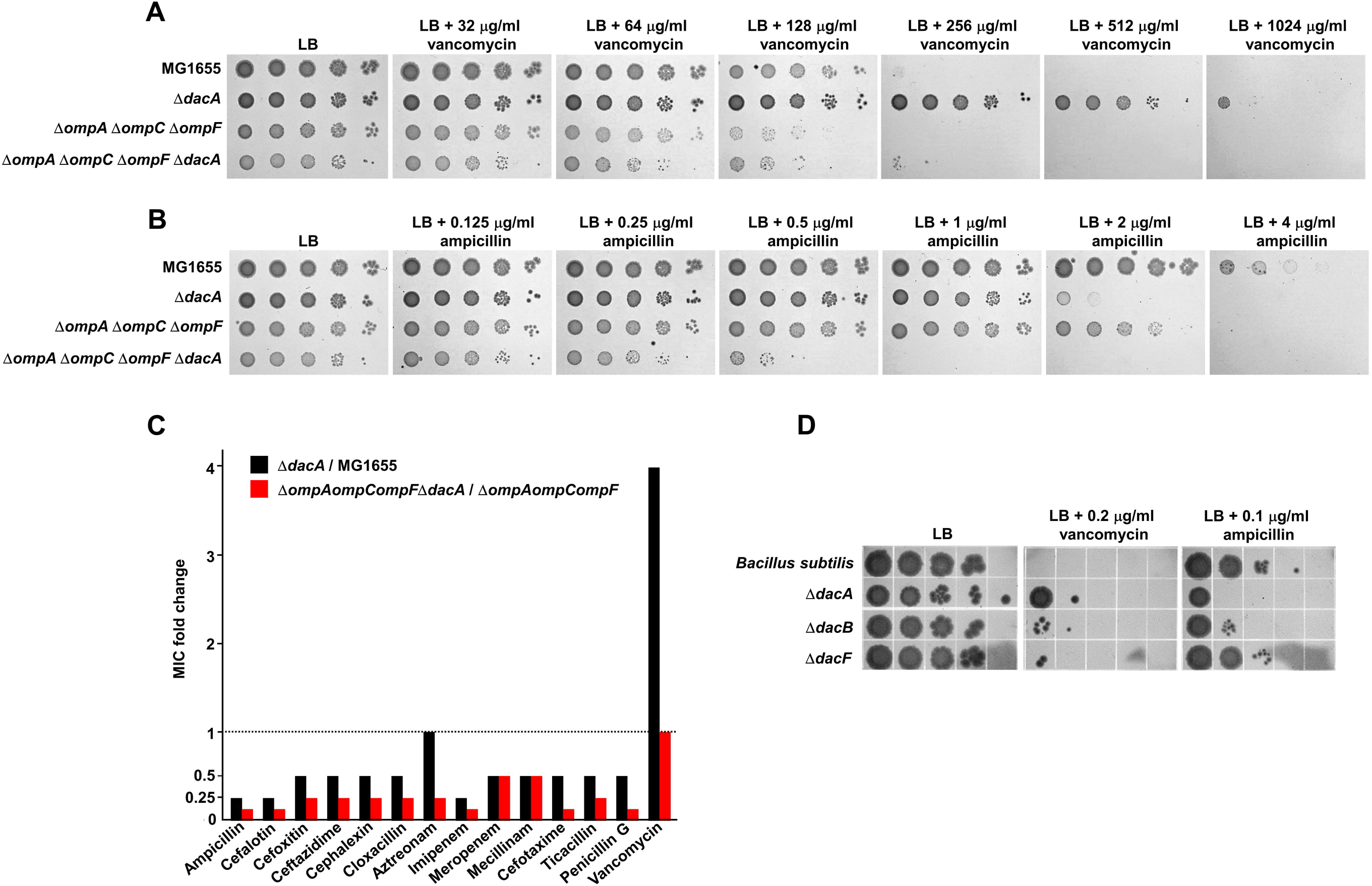
Effects of the OM permeability barrier on vancomycin resistance and β-lactam sensitivity in the *dacA* mutant. (A) The effect of the OM permeability barrier on vancomycin resistance. The wild-type and indicated mutant cells were serially diluted from 10^8^ to 10^4^ cells/ml in 10-fold steps and spotted onto an LB plate and LB plates containing the indicated vancomycin concentrations. (B) The effect of the OM permeability barrier on ampicillin sensitivity. The wild-type and indicated mutant cells were serially diluted from 10^8^ to 10^4^ cells/ml in 10-fold steps and spotted onto an LB plate and LB plates containing the indicated ampicillin concentrations. (C) The effect of the OM permeability barrier on the MICs of β-lactams. The MICs of various antibiotics were measured against the indicated mutant strains. The relative MIC values for the *dacA* mutant cells compared to those for the wild-type cell are shown as black bars, while the relative MIC values for the *ompA ompC ompF dacA* mutant cells compared to those for the *ompA ompC ompF* mutant cells are shown as red bars. (D) Vancomycin resistance and β-lactam sensitivity of the *B. subtilis dacA* mutant. The wild-type and indicated mutant cells were serially diluted from 10^8^ to 10^4^ cells/ml in 10-fold steps and spotted onto an LB plate and LB plates containing the indicated antibiotic concentrations.

### The divergent effect of PG carboxypeptidase on ampicillin and vancomycin susceptibility is also detected in *Bacillus subtilis*

We also examined the effect of PG carboxypeptidase on vancomycin and ampicillin susceptibility in *B. subtilis*, a Gram-positive bacterium that lacks the OM. *B. subtilis* has three homologs of *E. coli* DacA (DacA, DacB, and DacF), and the *dacA* mutant of *B. subtilis* was resistant to vancomycin, while the *dacA* and *dacB* mutants were sensitive to ampicillin (Fig. 7D). Both vancomycin resistance and ampicillin sensitivity of *B. subtilis dacA* mutant were significantly weaker than those of *E. coli dacA* mutant. Vancomycin can easily access PG in the absence of the OM, which might have affected vancomycin resistance in the *B. subtilis dacA* mutant. Similarly, easy access of vancomycin to PG completely abolished vancomycin resistance in the *E. coli dacA* mutant (Fig. 7A), whereas the *B. subtilis dacA* mutant showed weak vancomycin resistance. This discrepancy may have been caused by the thick PG layer of *B. subtilis*, where the amount of the decoy D-Ala-D-Ala residues was relatively significantly higher than that in *E. coli*. The reason behind the relatively weak β-lactam sensitivity of the *dacA* mutant of *B. subtilis* remains unclear. Collectively, these results show that the divergent phenotypes of the *dacA* mutant for vancomycin and ampicillin susceptibility are also conserved in a Gram-positive bacterium, and the presence of the OM can influence the phenotype of the *dacA* mutant.

## DISCUSSION

PG is an essential architecture for bacterial growth and shape maintenance, and its biosynthesis is inhibited by many antibiotics, such as β-lactams and vancomycin, leading to cell lysis. Although extensive studies have been conducted on PBPs, the physiological roles of PG hydrolases are poorly understood, especially regarding their impacts on antibiotic resistance. In this study, we revealed an intriguing connection between PG carboxypeptidases and antibiotic resistance. The PG carboxypeptidase-defective mutant showed β-lactam sensitivity and vancomycin resistance. Based on various genetic and biochemical experiments, we proposed the molecular mechanisms underlying these divergent phenotypes. Vancomycin resistance was caused by a decoy mechanism associated with increased D-Ala-D-Ala residues. Meanwhile, β-lactam sensitivity was associated with the physical interactions between DacA and PBPs, and these interactions required the C-terminal membrane-anchoring domain of DacA. Both mechanisms were independent of LD-transpeptidases. Intriguingly, increased permeability of the OM strongly weakened the vancomycin resistance of the *dacA* mutant, whereas it significantly strengthened β-lactam susceptibility. Collectively, our results demonstrate two distinct molecular mechanisms of PG carboxypeptidase-mediated antibiotic resistance in *E. coli*, a Gram-negative bacterium.

*Staphylococcus aureus*, a Gram-positive bacterium, has one orthologue of DacA which is known as PBP4 (hereinafter referred to as SaPBP4). SaPBP4 displays 25% identity (45% similarity) with *E. coli* DacA, and these two proteins have a similar molecular weight. Many clinical isolates with decreased expression level of SaPBP4 showed vancomycin resistance or β-lactam sensitivity (32–36). Previous studies suggested that the decoy mechanism of D-Ala-D-Ala is associated with vancomycin resistance (33, 35), which is consistent with the results of this study. However, unexpectedly, many recent reports based on *in vivo* and *in vitro* data showed that SaPBP4 has a strong DD-transpeptidase activity, but a weak DD-carboxypeptidase activity (37–40). Because many *in vitro* and *in vivo* experiments have demonstrated DD-carboxypeptidase activity of *E. coli* DacA (19, 20, 41), a direct comparison between our study and SaPBP4-related studies seems to be difficult. However, our study demonstrated that DacA physically interacted with DD-transpeptidases, namely PBPs, and these interactions were strongly associated with the effect of DacA on β-lactam resistance. This implies that *E. coli* DacA also affects DD-transpeptidases, that is, the formation of 4-3 cross-links. Because the two phenotypes of the *dacA* mutant identified in this study did not have a strong association with LD-transpeptidases, the DD-transpeptidase-related function of PG carboxypeptidases should be investigated in *E. coli* in the further studies. Notably, a study demonstrated that SaPBP4 can exert β-lactamase activity, based on *in vitro* data using nitrocefin (42). However, the β-lactamase activity of *E. coli* DacA has not been reported. In summary, these results suggest that further studies are required to elucidate the primary physiological role of PG carboxypeptidase in both Gram-negative and Gram-positive bacteria.

The mechanism of vancomycin resistance has been poorly analyzed in Gram-negative bacteria, including *E. coli*, as they are intrinsically resistant to vancomycin due to the presence of OM. Our study revealed decoy-mediated mechanism of vancomycin resistance in *E. coli*, indicating that there are various molecular mechanisms of vancomycin resistance in *E. coli*, besides the inhibition of vancomycin passage by the OM barrier. Although the decoy-mediated mechanism of vancomycin resistance was firstly suggested in *S. aureus* based on SaPBP4-related studies (42), the decoy model in *S. aureus* should be modified because SaPBP4 is a DD-transpeptidase, not a DD-carboxypeptidase. In this study, we clearly showed through an in vitro experiment, using purified PG (Fig. 4A), and an in vivo experiment, using DacA(ΔC) mutant (Fig. 5B), that vancomycin resistance in the *dacA* mutant is induced by decoy-mediated mechanism. Therefore, our study firstly revealed decoy-mediated vancomycin resistance mechanism of PG carboxypeptidase, at least in *E. coli*. Molecular decoy-mediated vancomycin resistance in *E. coli* has also been reported in a strain with a point-mutation in the *waaL* gene (encoding O-antigen ligase), which results in the synthesis of PG-modified lipopolysaccharide (LPS) (43). PG-modified LPS was transported through the OM and induced the decoy-mediated vancomycin resistance (43). These results imply that further studies are required to identify the diverse mechanisms associated with vancomycin resistance in *E. coli*.

The relationship between PG carboxypeptidase and β-lactam resistance has been consistently studied in *E. coli* (2, 13–15). The β-lactam sensitivity of the *dacA* mutant can be induced by diverse probable mechanisms as follows: dependency on 4-3 cross-links, β-lactamase activity of DacA, and the decoy model by class C PBPs, including DacA. Unexpectedly, our data showed that all the suggested mechanisms were not associated with the β-lactam sensitivity of the *dacA* mutant. Instead, our results suggest that the physical interactions of DacA with PBPs are strongly associated with the β-lactam sensitivity of the *dacA* mutant. It is necessary to determine whether this conclusion can be applied to other Gram-negative bacteria. The physiological significance of the physical interactions of DacA with PBPs is currently unknown. However, these results strongly imply that DacA plays diverse physiological roles, beyond the regulation of 3-3 cross link formation and Lpp-PG attachment.

The OM acts as a strong barrier against the influx of antibiotics owing to the hydrophobic acyl chains of LPS and hydrophilic lateral interactions between LPSs bridged by divalent cations (44, 45). Non-specific porins strongly regulate OM permeability (31). Our study again demonstrated that the presence of the OM strongly affected antibiotic resistance of Gram-negative bacteria. The vancomycin resistance of the *dacA* mutant was almost completely abolished in the cells with increased OM permeability, while the β-lactam sensitivity of the *dacA* mutant was significantly increased under the same condition (Fig. 7). These opposing patterns imply that the effect of OM permeability barrier is diverse and complex for the different types of antibiotics.

Although the C-terminal membrane-anchoring domain of DacA is known to be important for its function in morphology and its localization at the cell division site (16, 30), the underlying molecular mechanism has not been elucidated. Our data showed that the C-terminal domain of DacA was necessary for its interactions with PBPs, including PBP1a, PBP1b, PBP2, and PBP3 (Fig. 6). This result successfully explains why the C-terminal domain of DacA is required for its function in morphology and its localization at the septum. PBP2 and PBP3 play pivotal roles in cell morphology and their inhibitions cause cell rounding and filamentation, respectively (46). PBP3 is localized to the septum and forms the divisome complex with various proteins involved in cytokinesis (1, 47). Therefore, the role of the C-terminal domain in morphology might be associated with the physical interactions with PBP2 and PBP3, and its role in localization at the septum might be related to the physical interaction with PBP3. These findings should be confirmed by conducting further studies. Beause the DacA(ΔC) mutant without its C-terminal domain could not complement all the phenotypes of the *dacA* mutant, besides vancomycin resistance, the main function of DacA is associated with its C-terminal domain, that is, the interactions with PBPs. To complement the β-lactam sensitivity of the *dacA* mutant, the interactions with PBPs as well as the enzymatic activity of DacA is required (Fig. 2B). This implies that DacA does not affect PBPs only physically. The exact physiological significance of the DacA-PBP complexes remains unclear at present, and thus, further studies are required.

Our study showed the inverse phenotypes of the PG carboxypeptidase-defective mutant for vancomycin and β-lactam resistance and revealed distinct molecular mechanisms responsible for these phenotypes. Therefore, we provide novel insights into the roles of PG carboxypeptidases in the regulation of antibiotic resistance. Our findings also provide future research directions to elucidate the LD-transpeptidase-independent physiological function of PG carboxypeptidase.

## MATERIALS AND METHODS

### Bacterial strains, plasmids, and culture conditions

All the strains and primer sequences used in this study are presented in Table S1 and Table S2, respectively. All *E. coli* deletion strains were constructed using λ red recombinase, as previously described (48), with some modifications. An FRT-flanked kanamycin resistance gene in a plasmid pKD3 was amplified by conducting polymerase chain reaction (PCR) using deletion primer sets (See Table S2) with 50-mer sequences for the recombination between each target gene and the kanamycin gene. The PCR products were purified using PCR purification kit (Qiagen, USA) and transformed into MG1655 cells harboring a plasmid pKD46 that expresses λ red recombinase. After regeneration at 37°C, the cells were spread onto Luria-Bertani (LB) plates containing kanamycin (50 μg/ml) and incubated overnight at 37°C. The gene deletion in kanamycin-resistant cells was confirmed by PCR using deletion-confirming primer sets. To remove the kanamycin resistance gene, a plasmid pCP20, which expresses FLP recombinase, was transformed into deletion strains. After incubation at 37°C, the recombination between FRT sequences was checked by PCR using deletion-confirming primer sets, and the curing of the plasmid pCP20 was checked by growth failure in an LB plate containing ampicillin (100 μg/ml). To minimize the physiological change, the curing of the plasmid pCP20 was performed at 37°C, as previously described (49). All transformation procedures were performed by electroporation. Additional gene deletion, such as generation of the *dacA dacD* double mutant, was also carried out using λ red recombinase. Experimental procedures, including pKD46 transformation, gene disruption, pCP20 transformation, and FRT recombination, were sequentially performed. The *B. subtilis* deletion mutant strain was constructed using long-flanking homology PCR as previously described (50, 51).

The MG1655 PBP1a-Flag strain was also constructed using λ red recombinase, as previously described (49). Briefly, the DNA sequence including the 3 × Flag gene and the chloramphenicol resistance gene was amplified from the plasmid pBAD-PBP1a-Flag. After overnight DpnI digestion for the removal of template plasmids, PCR products were transformed into the MG1655 strain containing pKD46. The chromosomal fusion of the 3 × Flag tag to the C-terminus of PBP1a was confirmed by PCR. Other chromosomal PBP-Flag strains were also constructed through the same method. The MG1655 DacA(ΔC)-Flag strain was constructed through some modified process to remove the antibiotic resistance gene. The FRT-flanked kanamycin resistance gene in the plasmid pKD3 was inserted into the plasmid pBAD-Flag, generating the plasmid pBAD-Flag-FRT-Kan. The DNA sequence including the *3 × Flag* gene and the kanamycin resistance gene was amplified from the plasmid pBAD-Flag-FRT-Kan. After the chromosomal fusion of the 3 × Flag tag to the 385^th^ amino acid residue of DacA, the kanamycin resistance gene was removed by the recombination between FRT sequences using the plasmid pCP20, as described above.

The pBAD24a-based expression vectors of PG carboxypeptidases were constructed using In-Fusion cloning (Clontech, USA). The entire open reading frame of PG carboxypeptidase genes was amplified by PCR using cloning primer sets (See Table S2) containing the 15-mer sequence at the 5’ end for recombination with the vector. The PCR products were purified using PCR purification kit (Qiagen, USA) and *in vitro* recombination was performed between purified PCR products and pBAD24a plasmids digested by NdeI and BamHI restriction enzymes. Plasmid construction was confirmed by sequencing analysis. Through a similar method, the pRE1-based expression vector of DacA was constructed. Because the promoter in the pRE1 plasmid is recognized by *E. coli* RNA polymerase, but not T7 RNA polymerase, and is under the control of the cI protein of phage λ, the gene in the pRE1 plasmid is constitutively expressed in the general *E. coli* cells lacking the cI protein, including MG1655 (52, 53). To overexpress DacA, the pET28a-based or pET24a-based vector expressing His-tagged DacA or non-tagged DacA, respectively, was constructed through a similar method. In these cases, a truncated *dacA* gene without the DNA sequence encoding the signal sequence (1-29 amino acid residues) was cloned into the pET-based plasmids. The pRE1-DacA(S73G) and pRE1-DacA(K242R) plasmids were constructed by PCR using the pRE1-DacA plasmid as a template and DpnI digestion, as previously described (6).

### Determination of MIC values

The MIC values of the antibiotics were determined according to the Clinical Laboratory Standards Institute guideline (54). Cells from overnight seed culture grown in LB medium were inoculated into Mueller-Hinton broth and cultured at 37°C to a McFarland turbidity standard of 0.5 (approximately 1.5 × 10^8^ cells/ml). Cultured cells were diluted with Mueller-Hinton broth to reach a final concentration of 10^7^ cells/ml. Dilution samples (10 μl) were spotted onto Mueller-Hinton plates containing antibiotics of final concentrations of 512 μg/ml to 7.8 ng/ml in two-fold serial dilutions, and the plates were incubated at 37°C for 20 h. The MIC value of each antibiotic is defined as the lowest concentration of antibiotic at which the lawn growth of cells is inhibited.

### Purification of PG and *in vitro* binding assay with fluorescent vancomycin

The cells (1 L) of the wild-type and *dacA* mutant strains grown in LB medium to early exponential phase were harvested and resuspended in 20 ml of phosphate buffered saline (PBS) buffer. After adding 80 ml of 5% SDS buffer, the mixtures were boiled at 100°C for 30 min and incubated at room temperature overnight. The samples were ultracentrifuged at 160,000 × *g* for 1 h at room temperature, using Himac Micro Ultracentrifuge CS120FNX (Hitachi, Japan). To remove the remaining SDS, the pellets were washed with distilled water (DW) at least five times. The washed pellets were resuspended in 1 ml of PBS buffer. To remove high-molecular-weight glycogen and proteins, α-amylase (final concentration of 0.2 mg/ml) and α-chymotrypsin (final concentration of 300 μg/ml) were added to the resuspended samples and the mixtures were incubated at 37°C overnight. After adding SDS (final concentration of 1%), the samples were boiled at 100°C for 30 min and ultracentrifuged at 260,000 × *g* for 15 min at room temperature. To remove the remaining SDS, the pellets were washed with DW three times. The washed pellets were resuspended in 1 ml of DW and the samples were stored at −80°C.

To perform *in vitro* binding assay with fluorescent vancomycin,, 50 mg of PGs purified from the wild-type and *dacA* mutant cells were mixed with 10 μg of BODIP™ FL vancomycin (Invitrogen, USA). After incubation at 37°C for 30 min, the mixtures were ultracentrifuged at 260,000 × *g* for 15 min at room temperature. After washing the pellets with DW once, the pellets were resuspended in 50 μl DW and the amount of bound fluorescent vancomycin was determined by measuring the fluorescence emission of fluorescent vancomycin at 513 nm following excitation at 490 nm.

### Purification of overexpressed proteins and β-lactamase assay using nitrocefin

ER2566 cells harboring the plasmid pET28a-DacA or pET28a-NDM-1 from overnight seed culture were inoculated into LB medium containing kanamycin (50 μg/ml). When the OD600 reached approximately 0.5, 1 mM isopropyl-β-D-1-thiogalactopyranoside (IPTG) was added into the culture medium. After overnight culture at 16°C, harvested cells were resuspended in buffer A [50 mM Tris-HCl (pH 8.0) and 150 mM NaCl] and disrupted by a French press cell at 10,000 psi. Soluble His-tagged DacA and NDM-1 proteins were separated through centrifugation at 8,000 × *g* for 20 min at 4°C. The supernatants containing soluble target proteins were loaded into a Talon metal affinity resin (Clontech, USA). After washing three times with buffer A, His-tagged proteins bound to Talon resins were eluted with buffer A containing 200 mM imidazole. After overnight dialysis with buffer B [50 mM Tris-HCl (pH 7.5) and 50 mM NaCl] at 4°C, purified proteins were promptly used for measuring the β-lactamase activity.

To measure β-lactamase activity, purified NDM-1 (11 μg) and DacA (160 μg) proteins were mixed with 1 mM nitrocefin in 100 mM MES (pH 7.0) buffer containing 1 mg/ml bovine serum albumin. The degradation of nitrocefin was detected by measuring the absorbance at 486 nm over time. Instead of purified protein, the sample containing a Tris buffer of the same volume was used as a control.

### Binding assay of Bocillin FL to PBPs

Cells (20 ml) of the wild-type and *dacA* mutant strains grown in LB medium to early exponential phase were harvested. After washing once with 1 ml PBS buffer, the cells were resuspended in 1 ml PBS buffer and disrupted by a French press cell. Crude extracts were mixed with 5 μg of Bocillin FL in the presence or absence of 8 μg ampicillin. After incubation at room temperature for 10 min, 100 μl of the samples was mixed with 100 μl of 5× SDS sample buffer, followed by boiling at 100°C for 5 min. The mixture (20 μl) was loaded onto 4–20% gradient Tris-glycine polyacrylamide gel and run at 180 V for 80 min. PBP proteins bound to Bocillin FL were detected by scanning the fluorescence emission at 511 nm following excitation at 504 nm.

### Assessment of the enzymatic activity of PG carboxypeptidases

The enzyme activity of purified PG carboxypeptidases was measured using the substrate diacetyl-L-Lys-D-Ala-D-Ala (AcLAA) (20). D-Ala was released by PG carboxypeptidase, and the released D-Ala was degraded into pyruvate, hydrogen peroxide, and amine group by D-amino acid oxidase (Sigma Aldrich, United States). Horseradish peroxidase (Sigma Aldrich, United States) reduces hydrogen peroxide to H_2_O using Amplex Red (Invitrogen, United States) as an electron donor, which in turn is converted into resorufin by oxidation. The level of resorufin can be measured at 563 nm spectrophotometrically. Purified proteins (1 μM) were incubated at 37°C with 50 mM Tris-HCl (pH 8.0) containing 1 mM AcLAA in final volume of 200 μl. After incubation for 60 min, the reaction was stopped by boiling at 100°C for 20 min. After centrifugation at 13,000 rpm for 5 min, only the supernatant was mixed with 800 μl of assay buffer with 50 mM HEPES-NaOH (pH 7.5), 10 mM MgCl_2_, 50 μM Amplex red, 54 μg/ml horseradish peroxidase, and 75 μg/ml D-amino acid oxidase. The mixed samples were incubated at 37°C for 60 min, and the absorbance at 563 nm was measured using UV-Vis spectrophotometer UVmini-1240 (Shimadzu, Japan).

### Assessment of physical interactions between DacA/DacA(ΔC) and PBPs

ER2566 cells harboring pET-based plasmid expressing His-tagged DacA or non-tagged DacA were cultured in 50 ml of LB medium at 37°C. When the OD600 was approximately 0.5, 1 mM IPTG was added and the cells were cultured at 30°C for 4 h. MG1655 chromosomal PBP-Flag strains were cultured in 100 ml of LB medium at 37°C to early stationary phase. Harvested ER2566 and MG1655 cells were resuspended in buffer A containing 1% sodium n-dodecyl-β-D-maltoside and disrupted by a French press cell at 10,000 psi. After centrifugation at 2,100 × *g* for 5 min at 4°C, the supernatants were mixed with 100 μl of the Talon metal affinity resin (Clontech, USA) in a 1.5-ml tube. After washing with 1 ml of buffer A containing 1% sodium n-dodecyl-β-D-maltoside three times, bound proteins were eluted using buffer A containing 200 mM imidazole. Eluted proteins were separated on 4–20% gradient Tris-glycine polyacrylamide gels (KOMA biotech, Korea) and were transferred onto PVDF membranes. The protein levels of PBPs were determined using monoclonal antibody against Flag-tag (Santa Cruz Biotechnology, United States), according to the standard procedure. A similar pull-down experiment was also performed using overexpressed full-length PBP1a or PBP1b and overexpressed His-tagged DacA or DacA(ΔC). Eluted proteins were separated on 4–20% gradient Tris-glycine polyacrylamide gels.

## ACKNOWLEDGMENTS

This work was supported by research grants from Basic Science Research Program through the National Research Foundation of Korea funded by the Ministry of Education (NRF-2020R1I1A2058026, 2021R1A6A3A01086677, and 2021R1A6A3A01086629).

